# Comparative transcriptome analysis by deep RNA sequencing at early stage of skin pigmentation in goats (*Capra hircus*)

**DOI:** 10.1101/079871

**Authors:** Hangxing Ren, Gaofu Wang, Jing Jiang, Liangjia Liu, Nianfu Li, Jie Li, Lin Fu, Haiyan Zhang, Risu Na, Yongfu Huang, Li Zhang, Lei Chen, Yong Huang, Peng Zhou

## Abstract

Although specific genes have been found to be associated with skin pigmentation, the global gene expression profile for the early stage of skin pigmentation and development in mammals is still not well understood. Here we reported a rare natural group of goat (Youzhou dark goat) featuring the dark skin of body including the visible mucous membranes, which may be an exclusive kind of large mammalian species with this special phenotype so far. In the present study, we characterized the 100-day-old fetal skin transcriptome in hyperpigmented (dark-skinned) and wild-type (white-skinned) goats using deep RNA-sequencing. A total of 923,013,870 raw reads from 6 libraries were obtained, and a large number of alternative splicing events were identified in the transcriptome of fetal skin, including the well-known melanogenic genes *ASIP*, *TYRP1*, and *DCT*, which were differentially expressed in the skin between the dark-skinned and white-skinned goats. Further analysis demonstrated that differential genes including *ASIP*, *TYRP1*, *DCT*, *WNT2*, *RAB27A*, *FZD4*, and *CREB3L1* were significantly overrepresented in the *melanogenesis* pathway and several biological process associated with pigmentation. On the other hand, we identified 1616 novel transcripts in goat skin based on the characteristics of their expression level and gene composition. These novel transcripts may represent two distinct groups of nucleic acid molecules. Our findings contribute to the understanding of the characteristics of global gene expression at early stages of skin pigmentation and development, as well as describe an animal model for human diseases associated with pigmentation.

## INTRODUCTION

Skin pigmentation is a complex process that includes melanin biosynthesis in melanocytes, which is transferred to keratinocytes (Wolff 1973; Seiberg 2001). Unlike the numerous variations of coat color, there are only a few natural variants of skin color in other mammals compared to the phenotypes of human skin color. As a result, most studies on the genetics of skin pigmentation are conducted in humans (Baxter and Pavan; Sturm 2009; Quillen and Shriver 2011; Meng *et al.* 2012) and mice (Bennett and Lamoreux 2003; Garcia *et al.* 2008), in which 378 color genes (171 cloned genes and 207 uncloned genes) have been identified (http://www.espcr.org/micemut) to date. However, only a small portion of these color genes have been identified as candidate causative genes for skin pigmentation in different populations of humans, mice/rats, and sheep, including *ASIP*, *TYRP1*, *DCT*, *TYR*, *HERC2*, *OCA2*, *MC1R*, *SLC24A5*, *SLC45A2*, *IRF4*, *KITLG* (Slominski *et al.* 2004; Yamaguchi *et al.* 2007; Ebanks *et al.* 2009; Yamaguchi and Hearing 2009; Garcia-Gamez *et al.* 2011; Kondo and Hearing 2011; Liu *et al.* 2013; Raadsma *et al.* 2013; Han *et al.* 2015). In addition, few studies have characterized the global gene expression profile of skin pigmentation and development. Although investigations of the skin transcriptome have recently been conducted on coat color (Fan *et al.* 2013) and hair follicles (Xu *et al.* 2013a; Xu *et al.* 2013b; Wang *et al.* 2015; Yue *et al.* 2015; Gao *et al.* 2016) in sheep and goats, and skin color in the common carp (Jiang and Bikle 2014b; Wang *et al.* 2014), red tilapia (Zhu *et al.* 2016), and chickens (Zhang *et al.* 2015), the genetic basis for skin pigmentation is not well understood compared to that of hair or coat color.

Here, we report a rare indigenous goat breed (Youzhou dark goat) that features dark skin of the body including the visible mucous membranes (Fig.1A), which is a rare fibromelanosis that has previously only been reported in the Silky fowl (Nozaki and Makita 1998; Muroya *et al.* 2000; Dorshorst *et al.* 2011; Shinomiya *et al.* 2012). The pigmentation phenotype of the Youzhou dark goat significantly differs from the piebald phenotype of the bovine (Weikard *et al.* 2013), whereas it is similar to a recently reported case of dermal melanocytosis in humans (Lee *et al.* 2010). However, the underlying mechanism of this hyperpigmentation is yet to be explored. Therefore, the Youzhou dark goat can be used as a medical model to study human diseases associated with pigmentation, such as skin melanopathy, melanosis coli, and mucosal melanosis. Interestingly, based on our long-term observations (unpublished), skin pigmentation parallels skin development during the pregnant and postnatal period in the Youzhou dark goat. Consequently, to understand fibromelanosis in mammals, it is necessary to investigate the biology of skin pigmentation at early developmental stages in goats. In developmental embryology, the growth of the fetal skin peaks at approximately 100 days of gestation in sheep and goats (Wang *et al.* 1996; Qin 2001). In the present study, we used the Youzhou dark goat (hyperpigmented or dark-skinned) and Yudong white goat (wild-type or white-skinned) as models of skin pigmentation to characterize the skin transcriptome in 100-day-old fetal goats using deep RNA-sequencing. Our study not only contributes to the understanding of the biology of skin pigmentation and development but also provides valuable information towards understanding human melanocytosis.

**Fig. 1.**
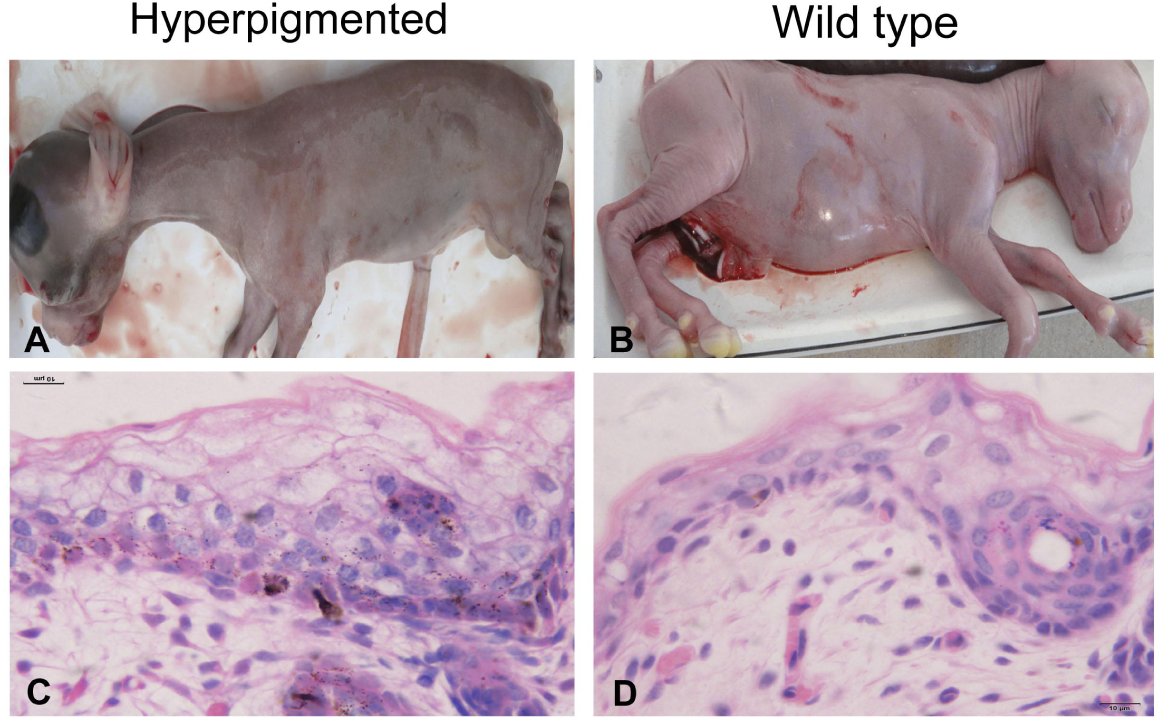
Histomorphological examination of the hyperpigmented and normal skin in fetal goats. To explore the histomorphological differences between the two breeds of goats at the early stage of skin pigmentation and development, the 100-day-old fetal skin from the Youzhou dark goat (A and C) and Yudong white goat (B and D) were examined using H & E staining. The scale bar for the images in (C) and (D) is 10 µm.

## MATERIALS AND METHODS

### Ethical statement

All surgical procedures in goats were performed according to the Regulations for the Administration of Affairs Concerning Experimental Animals (Ministry of Science and Technology, China; revised in June 2004) and adhered to the Reporting Guidelines for Randomized Controlled Trials in Livestock and Food Safety (REFLECT).

### Animals

Two goat groups with different skin pigmentation phenotypes were investigated in this study. The Yudong white goat (*Capra hircus*) is distributed in Southwest China (located at 31°14'-32°12' N and 108°15'-109°58' E) and features white color in the coat and skin. The Youzhou dark goat (*Capra hircus*) is an indigenous breed uniquely distributed in Youyang County in Chongqing, China (located at 26°54' N and 108°57' E) and features dark skin of the body including the visible mucous membranes but a generally white coat color. Briefly, three pregnant ewes from each breed were subjected to a caesarean section to collect the fetuses (n=3) at 100 days of gestation, and then the dorsal and ventral skins were collected from each fetus. The first sample (3 grams) was dissected and rapidly frozen whole in isopentane chilled over liquid nitrogen for histological examination. The second sample (3 grams) was snap-frozen in liquid nitrogen for RNA-sequencing and qPCR analysis.

### RNA isolation, library preparation and sequencing

In the present study, a total of 6 libraries were generated for sequencing according to the sample size (n=6) in two breeds of goats. For each of the 6 fetal goats, total RNA was isolated using TRIzol reagent (Invitrogen, USA) according to the manufacturer’s instructions. RNA degradation and contamination was monitored on 1% agarose gels. RNA purity was checked using the NanoPhotometer spectrophotometer (Implen, USA). RNA concentration was measured using a Qubit RNA Assay Kit with a Qubit 2.0 Flurometer (Life Technologies, USA). RNA integrity was assessed using the RNA Nano 6000 Assay Kit with the Agilent 2100 Bioanalyzer System (Agilent Technologies, USA). A total of 3 μg of RNA per sample was used as the input material for the RNA sample preparations. First, ribosomal RNA was removed using the Epicentre Ribo-zero™ rRNA Removal Kit (Epicentre, USA), and rRNA free residue was cleaned up using ethanol precipitation. Subsequently, the highly strand-specific libraries were generated using the rRNA-depleted RNA using the NEBNext Ultra™ Directional RNA Library Prep Kit for Illumina (NEB, USA) according to the manufacturer’s recommendations. Briefly, fragmentation was carried out using divalent cations under elevated temperature in NEBNext. First strand cDNA was synthesized using random hexamer primers and M-MuLV Reverse Transcriptase (RNaseH-). Second strand cDNA synthesis was subsequently performed using DNA Polymerase I and RNase H. In the reaction buffer, dNTPs with dTTP were replaced by dUTP. The remaining overhangs were converted into blunt ends via exonuclease/polymerase activities. After adenylation of the 3’ ends of the DNA fragments, NEBNext Adaptor with hairpin loop structure were ligated to prepare for hybridization. To select cDNA fragments of preferentially 150-200 bp in length, the library fragments were purified using the AMPure XP system (Beckman Coulter, USA). Then, 3 μl of USER Enzyme (NEB, USA) was used with size-selected, adaptor-ligated cDNA at 37°C for 15 min followed by 5 min at 95°C before PCR. PCR was performed with Phusion High-Fidelity DNA polymerase, Universal PCR primers and Index (X) Primer. Finally, the products were purified (AMPure XP system) and the library quality was assessed on the Agilent 2100 Bioanalyzer System. The clustering of the index-coded samples was performed on a cBot Cluster Generation System using TruSeq PE Cluster Kit v3-cBot-HS (Illumina) according to the manufacturer’s instructions. After cluster generation, the libraries were sequenced on an Illumina Hiseq 2000 platform and 100 bp paired-end reads were generated.

### Quality control

The raw data were firstly processed through in-house perl scripts. In this step, clean data were obtained by removing the reads containing adapters, reads containing over 10% of ploy-N sequences, and low quality reads (more than 50% of bases with Phred scores less than 5%) from the raw data. The Phred score (Q20, Q30) and GC content of the clean data were calculated. All the subsequent analyses were based on the high quality data. The sequencing data were submitted to the Genome Expression Omnibus (Accession Numbers GSE69812) in NCBI.

### Mapping to the reference genome

The reference genome and gene model annotation files were downloaded directly from the genome website (http://goat.kiz.ac.cn). The index of the reference genome was built using Bowtie v2.0.6 (Langmead *et al.* 2009; Langmead and Salzberg 2012) and paired-end clean reads were aligned to the reference genome using TopHat v2.0.9 (Trapnell *et al.* 2012; Kim *et al.* 2013).

### Transcriptome assembly

The goat reference genome and gene model annotation files were downloaded directly from the genome website (http://goat.kiz.ac.cn). The index of the reference genome was built using Bowtie v2.0.6 (Langmead *et al.* 2009; Langmead and Salzberg 2012) and paired-end clean reads were aligned to the reference genome using TopHat v2.0.9 (Trapnell *et al.* 2012; Kim *et al.* 2013). The mapped reads of each sample were assembled using both Scripture (beta2) (Guttman *et al.* 2010) and Cufflinks (v2.1.1) (Trapnell *et al.* 2010) in a reference-based approach. Scripture was run with default parameters. Cufflinks was run with ‘min-frags-per-transfrag=0’ and ‘--library-type fr-firststrand’, with all other parameters set as default.

### Quantification of gene expression level

Cuffdiff (v2.1.1) was used to calculate the FPKM (fragments per kb for a million reads) of both lncRNAs and coding genes in each sample (Trapnell *et al.* 2010). For biological replicates (n=3), transcripts or genes with a *P*-adjust<0.05 were considered differentially expressed between the two groups of goats (dark-skinned and white-skinned).

### Alternative splicing analysis

Alternative splicing (AS) events were classified into 12 basic types using the software Asprofile v1.0 (Florea *et al.* 2013). The number of AS events in each of the 6 samples was estimated separately. We used DEXSeq software(Anders *et al.* 2012) for the differential exon usage analysis of the AS transcripts, in which a general linear model was employed for the differential analysis of exon expression and a *P*-adjust<0.05 indicated a significant result.

### Identification of novel transcripts

To identify the novel transcripts from clean data, we first distinguished the novel transcripts and the candidate long noncoding RNAs using four tools including CNCI (v2) (Sun *et al.* 2013), CPC (0.9-r2) (Kong *et al.* 2007), Pfam-scan (v1.3) (Punta *et al.* 2012), and PhyloCSF (v20121028) (Lin *et al.* 2011). Transcripts predicted to be without coding potential by all of four tools above were filtered away (which are considered as the candidate long noncoding RNAs), and those with coding potential (which are selected by either of four tools above) but lacking of any known annotation were kept and considered as the novel transcripts in the present analysis. Quantification of gene expression level were estimated by calculating FPKMs of the novel transcripts.

RepeatMasker (http://www.repeatmasker.org/cgi-bin/WEBRepeatMasker) was used with the default parameters to identify various TE components in goat. To identify the position bias of TEs in the novel transcript, we searched the TEs in the 2, 000 bp upstream of TSS (Transcription Start Site) of each transcript identified in the goat genome (http://goat.kiz.ac.cn) and plotted read coverage at TSSs with the ggplot2 package in R(Wickham 2009).

### Differential expression and functional enrichment analysis

Cuffdiff provides statistical routines to determine differential expression in digital transcript or gene expression data using a model based on the negative binomial distribution (Trapnell *et al.* 2010). For biological replicates, transcripts or genes with a *P*-adjust<0.05 were considered differentially expressed. To identify the molecular events or cascades involved, the differentially expressed genes or lncRNA target genes were analyzed using the DAVID platform (Huang Da *et al.* 2009b; Huang Da *et al.* 2009a). Significance was expressed as a *P*-value, which was calculated using the EASE score (a *P*-value of 0.05 was considered significant).

### Validation of gene expression in RNA-seq using quantitative PCR

To validate the gene expression in RNA-seq, the total RNA from the RNA-seq analysis was used for qPCR (Fig. 4). Briefly, first strain cDNA was obtained using a One Step cDNA Synthesis Kit (Bio-Rad, USA), and the mRNA was then quantified using a standard SYBR with *GAPDH* as an endogenous control. Quantitative PCR was performed under the following conditions: 95°C for 30 sec, 40 cycles of 95°C for 5 sec, and the optimized annealing temperature for 30 sec. The primers and annealing temperatures for the 8 genes are listed in Table S1. All reactions were performed in triplicate for each sample. Gene expression was quantified relative to *GAPDH* expression using the 2^(-ΔCt)^ method. Corrections for multiple comparisons were performed using the Holm-Sidak method.

**Fig. 4.**
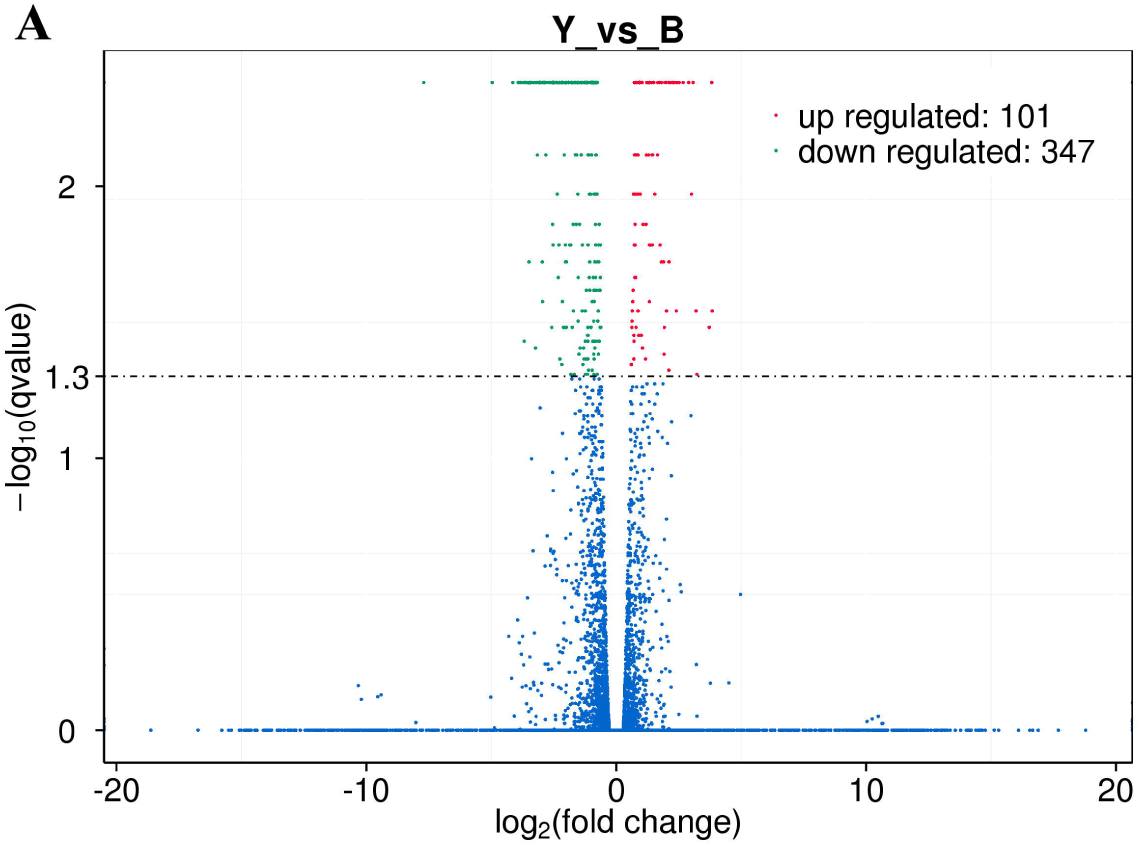
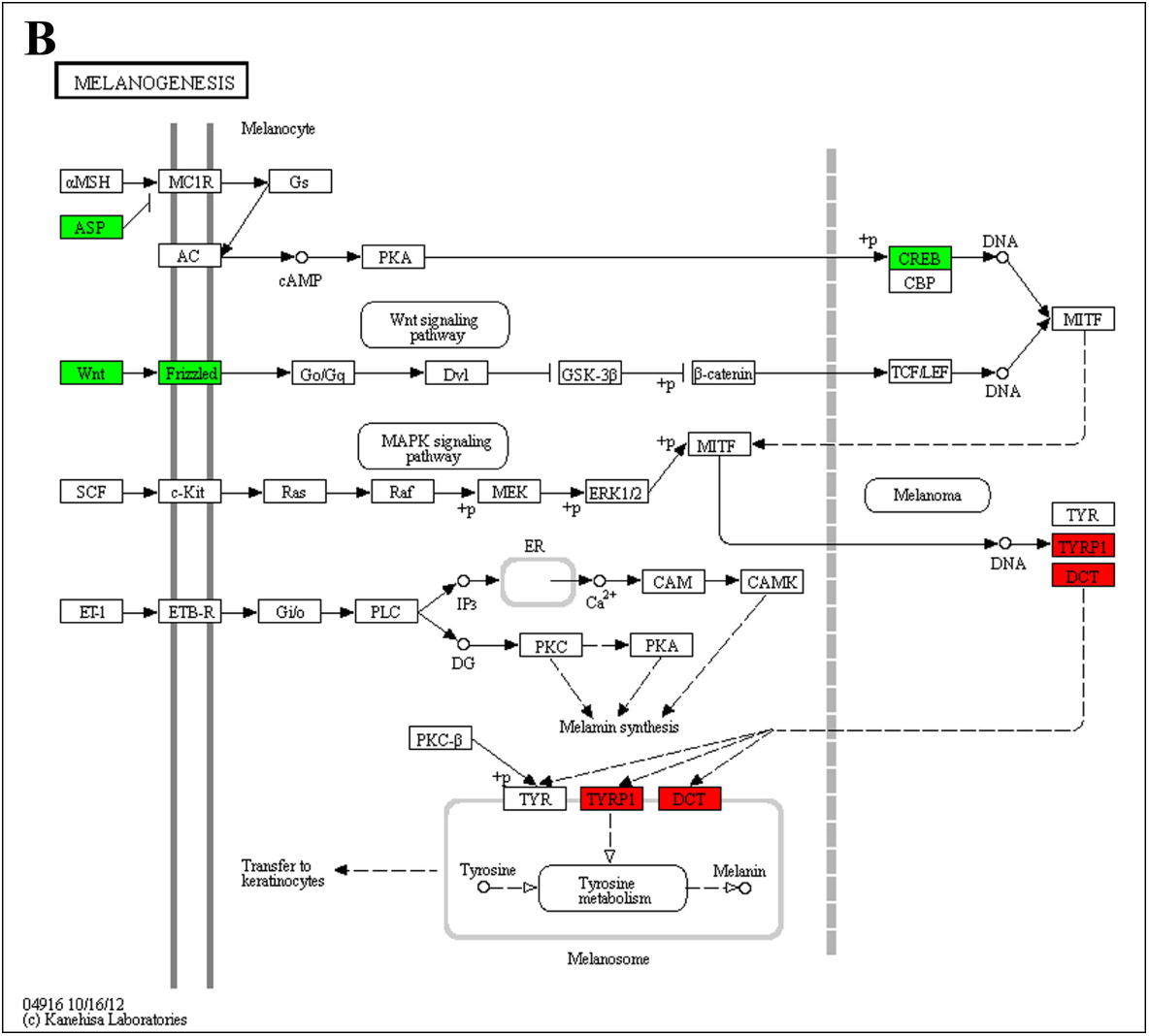
The differentially expressed genes between the dark- and white-skinned goats. (A) The differential expression of genes was identified based on the FPKM values (corrected *P*-value ≤ 0.05 and fold change ≥ 1.5) in the two groups. The volcano plot was generated based on the values above. Y indicates the dark-skinned goats, and B refers to the white-skinned goats. (B) Enrichment of the DE genes was conducted under KEGG pathways. Red indicates the up-regulated DE genes and green refers to the down-regulated genes in the dark-skinned goats compared to the control goats (white-skinned).

### Statistical analysis

Data analyses were performed using the R statistical package.

### Data availability

Figure S1 contains classification of the raw reads by RNA-seq for each of the libraries (samples) in the hyperpigmented and normal fetal goats. Figure S2 contains a heatmap of the cluster analysis for the differentially expressed genes. Table S1 contains statistics of the mapping reads to the reference genome. Table S2 contains annotation of novel transcripts in Swiss-Prot database. Table S3 contains differential exon usage (DEU) analysis of the mRNAs in goat fetal skin. Table S4 contains the differentially expressed genes between the wild-type and hyperpigmented goat skins. Table S5 contains results of GO and KEGG analysis of the up-regulated DE genes using DAVID. Table S6 contains results of functional annotation clustering of the DE genes using DAVID.

## RESULTS

### RNA sequencing of skin in fetal goats

As for phenotype, there are obvious differences in skin tissue pigmentation between the two breeds of fetal goat (Fig. 1A and B). Then we examined the phenotypic differences at the histological level using H & E staining. The results demonstrated that there were more melanin granules in the 100-day fetal epidermal and dermal layers in the Youzhou dark goat than in the Yudong white goat (Fig. 1C and D). To further explore the mechanism underpinning the differences in skin pigmentation, we characterized the skin transcriptome using RNA-seq performed with the Illumina HiSeq 2000 platform. In the present study, we obtained a total of 923,013,870 raw reads from 6 libraries (samples), of which 841,895,634 clean reads remained for further analysis after discarding the raw reads that contained adapter sequences, N sequences and low quality sequences. The percentage of clean reads among the raw tags in each library ranged from 88.39% to 93.02% (Fig. S1). We then mapped the clean reads to the goat reference genome (http://goat.kiz.ac.cn). Of the total reads from each library, more than 83% matched to a unique genomic location, whereas only 3% matched to multiple genomic locations (Table S1). The Phred score (Q20) for each library was greater than 95.4%, which indicates our RNA-seq data were high quality and suitable for the subsequent analyses.

### Novel transcripts in goat genome

We identified a total of 27,947 mRNAs and 1616 novel transcripts in fetal goat skin in the present study. To examine their homology with proteins in state-of-the-art database, we made a BLAST alignment for these novel transcripts in Swiss-Prot database (http://www.uniprot.org/). Results showed that 660 of them had a similarity in various degree, but with a relative lower coverage to the known proteins (Table S2). Another 956 novel transcripts were found without annotation in the state-of-the-art database presently. Then we examined the differences in gene length and expression levels (FPKMs) between the two groups of novel transcripts, and found that there were significant differences in gene length (Kolmogorov-Smirnov test, *P* = 0.011; Fig 2A) and expression levels (Kolmogorov-Smirnov test, *P* = 0.000; Fig 2B) between them. To further characterize the two groups of novel transcripts, we examined the composition of transposable elements (TEs) harboring in their gene sequences. Our results demonstrated that there were considerable differences in density of various TE components between the two groups of novel transcripts, as well as between the novel transcripts and mRNAs (Fig 2C). Further investigation of position bias for TEs relative to TSS (transcription start site) revealed that the LINE/RTE-BovB were higher enriched in novel transcripts without annotation than the other two groups (Fig 2D). These above findings suggest that the identified transcripts may represent different types of novel transcripts in goat genome.

**Fig. 2.**
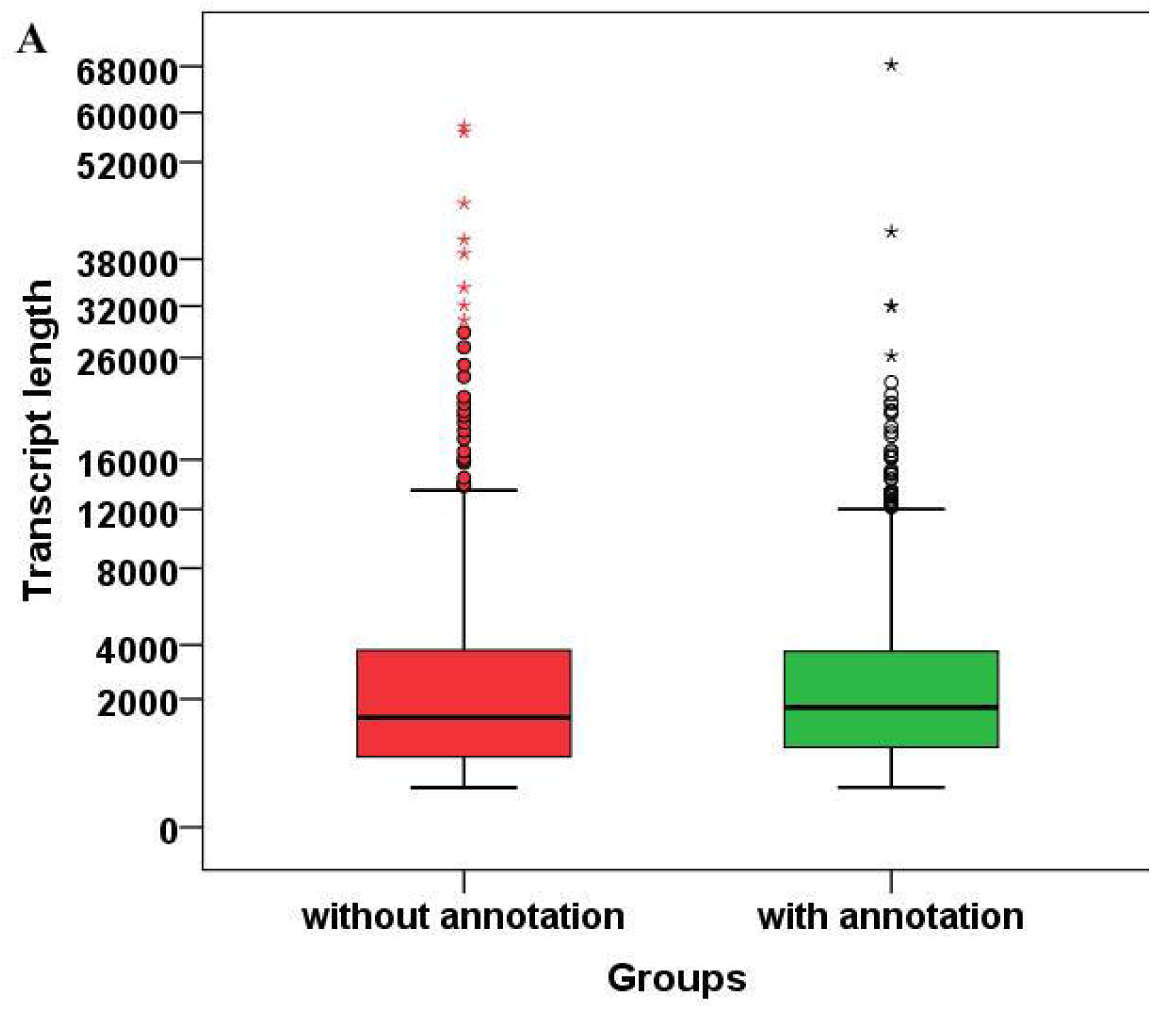
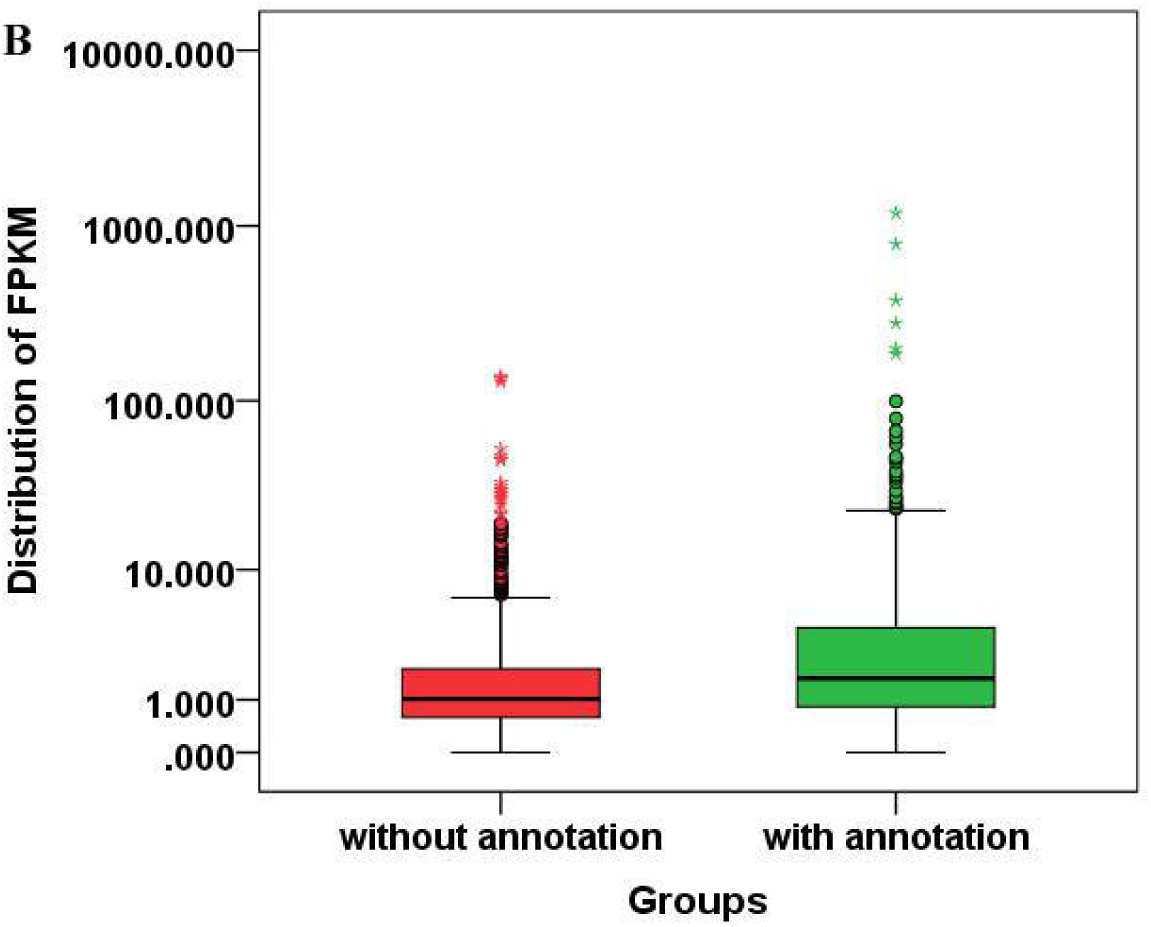
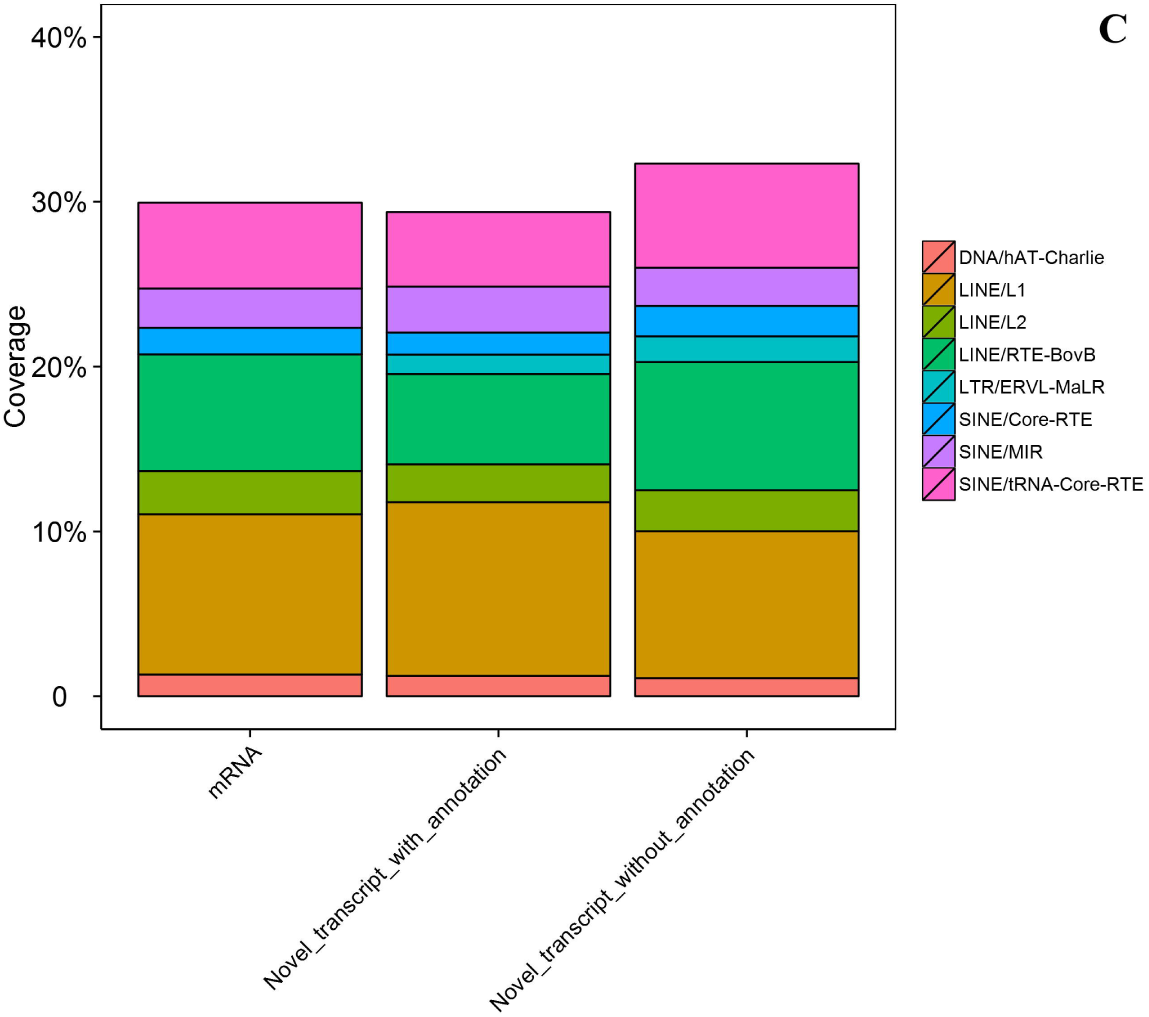
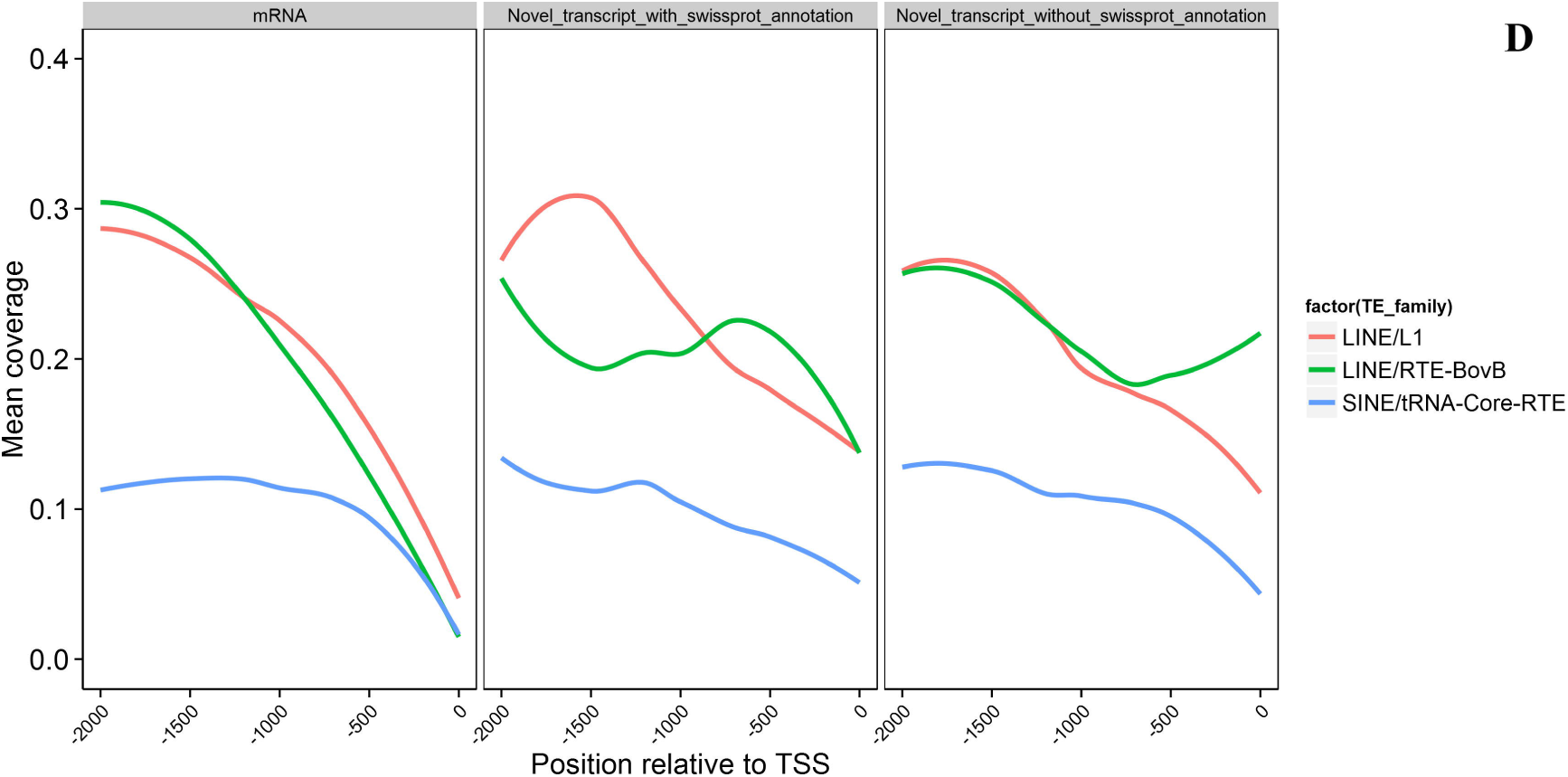
Characteristic differences between two groups of novel transcripts in goat genome. The transcript length (A) and expression level (B) of two classes of novel transcripts were compared using the Kolmogorov-Smirnov test, and a P value of 0.05 indicates significance between two groups. In two box plots, the circle indicates the outlier, and the asterisk labels the extreme. (C) The proportion of the main TE families in two groups of novel transcripts and mRNAs in the goat genome (http://goat.kiz.ac.cn). Differences in TE components between them were measured by using the Fisher Exact test. (D) The position bias of TE components in the 2,000 bp upstream of TSS in transcripts above mentioned.

### Alternative splicing of the fetal skin transcriptome

To ascertain the alternative splicing (AS) events of the skin transcriptome, we examined the data from six samples using ASprofile software. Our results showed that TSSs (alternative 5' first exon, transcription start sites), TTSs (alternative 3' last exon, transcription terminal sites), and SKIPs (skipped exons) were the three most frequently observed AS events among the 12 AS subtypes in each sample (Fig. 3A). Among the over 66,000 AS events, we observed AS events in two well-known genes involved in pigmentation, *ASIP* and *TYRP1*, which were found in both goat breeds. To further examine the AS events of the differentially expressed genes between the dark- and white-skinned goat breeds, we subsequently investigated differential exon usage of the AS events using DEXSeq software. The results demonstrated that 360 AS exons belonging to 253 known and 28 unknown genes were significantly differentially expressed between the dark- and white-skinned groups (*P*-adjust<0.05) (Table S3). Importantly, we found that some AS exons belonging to well-known genes involved in pigmentation, such as *ASIP*, *DCT*, and *RAB27A*, were significantly differentially expressed between the two different skin colors (Fig. 3B). This indicates that alternative splicing may be an important mechanism by which expression and function of ASIP, DCT, and RAB27A is regulated in melanocytes.

**Fig. 3.**
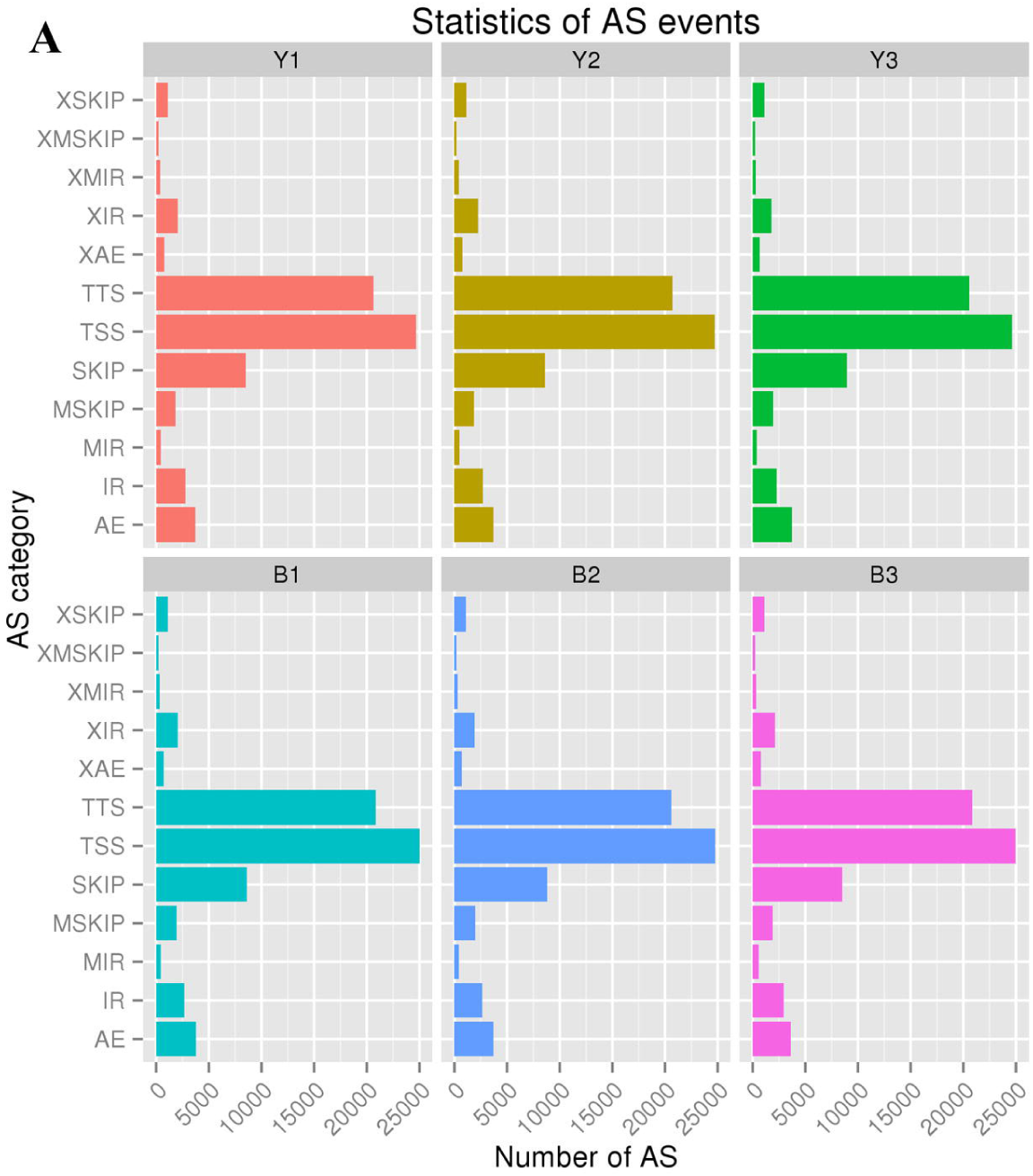
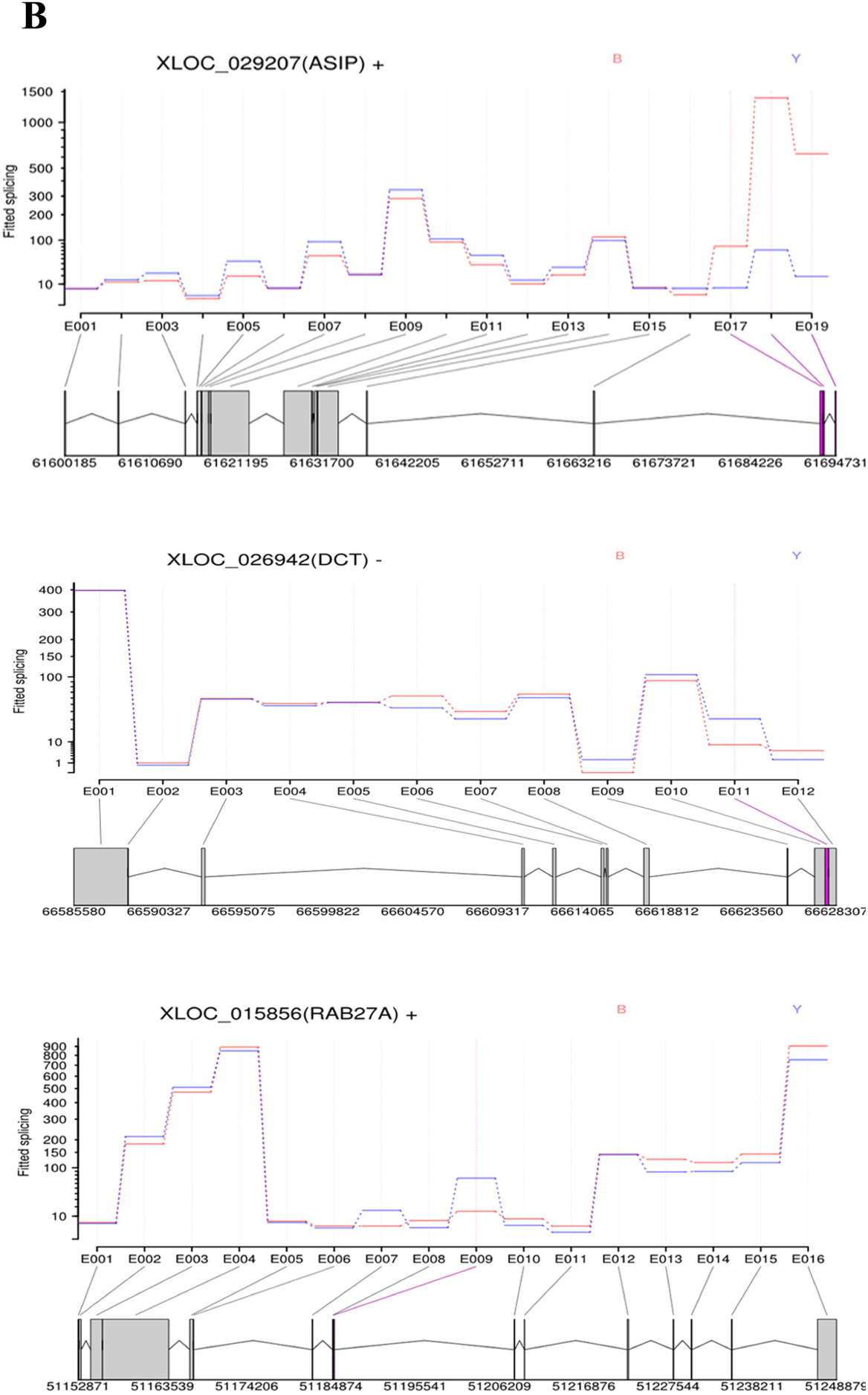
Alternative splicing (AS) events in the goat genome. Following the analysis using Cufflinks, the classification of AS events of transcripts were performed for each library (sample) using the ASprofile tool. The 12 main AS events (TSS, TTS, SKIP, XSKIP, MSKIP, XMSKIP, IR, XIR, MIR, XMIR, AE, and XAE) in the goat genome are summarized in (A). Y1, Y2, and Y3 indicate the dark skin goats, whereas B1, B2, and B3 indicate the white skin goats. (B) Differential exon usage of the AS events in the *ASIP* and *DCT* genes between the dark-skinned (Y, blue) and white-skinned (B, red) goats.

### Identification and functional clustering analysis of the differentially expressed genes

In our data, we obtained 448 differentially expressed (DE) genes (*P*-adjusted<0.05 and fold change≥1.5) using Cuffdiff between the dark- and white-skinned goats. Of the 448 DE genes, 101 genes were up-regulated and 347 were down-regulated in the dark-skinned goats compared with the white-skinned goats (Fig. 4A, Table S4, Fig. S2). Here, to elucidate the biology of skin pigmentation in goats, we specially focused on the up-regulated DE genes in the dark-skinned goats. Gene functional classification using DAVID demonstrated that a family of keratin (KRT) was significantly enriched in the up-regulated DE genes, including *KRT-I*, *KRT81*, *KRT83*, *KRTAP13-1*, *KRTAP11-1*, and *KRTAP7-1*, which are constitutive components of skin produced by keratinocytes. Gene ontology analysis showed that the top six biological processes were *response to inorganic substance*, *response to extracellular stimulus*, *response to reactive oxygen species*, *pigmentation during development*, *pigment biosynthetic process*, and *response to cAMP* (Table S5). KEGG analysis showed that genes involved in *melanogenesis* and the *MAPK signaling pathway* were significantly enriched. These results suggest that the up-regulated genes play important roles in the dark skin phenotype of the Youzhou dark goats.

To gain a comprehensive insight of the gene expression between the two breeds, we performed Gene ontology and pathway analyses of all DE genes between the normal and dark-skinned goats using DAVID. Functional clustering analysis of all DE genes showed that 65 annotation clusters were significantly enriched, particularly *cluster 41*, which comprises 7 terms related to hyperpigmentation including *pigment biosynthetic process*, *melanin biosynthetic process*, *pigment metabolic process*, *melanin metabolic process*, *secondary metabolic process*, *melanogenesis*, and *pigmentation during development*. Regarding the pathways involved, we identified 14 significantly overrepresented pathways (Table S6) including the *melanogenesis* pathway (Fig. 4B). Selected DE genes from the RNA-seq analysis were validated using quantitative PCR (Fig. 5). These findings provide additional evidence of the underlying diversity of dermal colors in goats on the transcriptome level. Moreover, we explored the enrichment of genes involved in human pigmentation diseases using DAVID to determine whether there are diseases with similar variations in dermal colors in goats as are observed in humans. As we expected, *ASIP* and *TYRP1* were significantly enriched in one type of OMIM disease (Table 1). This suggests that *ASIP* and *TYRP1* are the two likely candidate genes responsible for dermal hyperpigmentation in Youzhou dark goats.

**Fig. 5.**
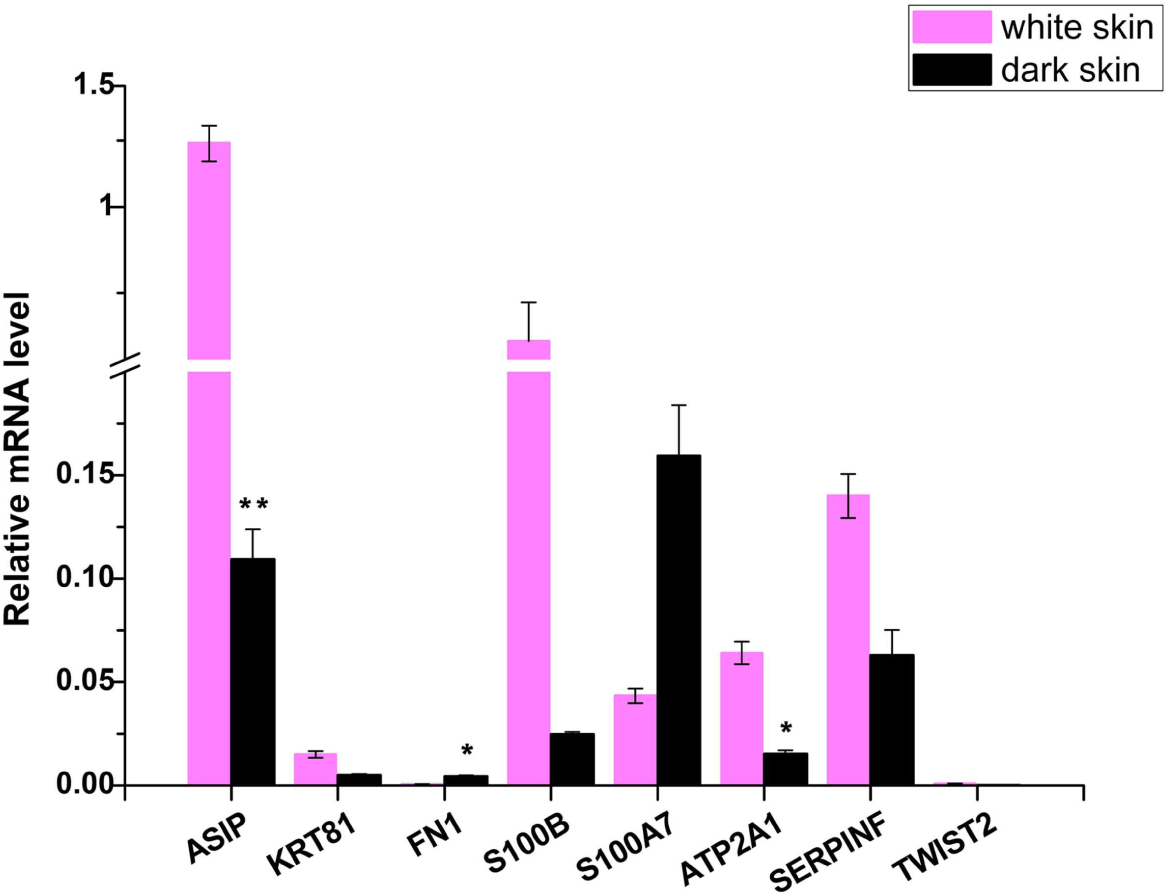
Validation of gene expression in the dark- and white-skinned goats using qPCR. Some of the identified melanogenic genes were examined in the dark- and white-skinned fetal goats using qPCR. Gene expression was quantified relative to *GAPDH* expression using the 2^(-ΔCt)^ method. Corrections for multiple comparisons were performed using the Holm-Sidak method. The data are shown as the mean ± 1 SE (n = 3). \**P*<0.05, \*\**P*<0.01.

## DISCUSSION

Although transcriptomic investigations on variations in skin color have been conducted in the common carp (Jiang and Bikle 2014b; Wang *et al.* 2014), red tilapia (Zhu *et al.* 2016), and chickens (Zhang *et al.* 2015), few studies have investigated this topic in mammals to date. We first characterized the skin transcriptome in two breeds of fetal goats that feature different skin colors using deep RNA-seq methods. The present study produced a large amount of data (over 84.2 Gb of clean reads for six samples) and more than 86% of the total reads were mapped to the goat reference genome (http://goat.kiz.ac.cn), which indicated that the RNA-seq data were of high quality and valuable for further analysis. First, we identified a significant number of AS events (Fig. 3A) in the skin transcriptome of fetal goats, which suggests that gene transcription and regulation is complex in the skin during this developmental stage in goats. AS events are tissue specific and widespread in the genome (Xu *et al.* 2002). AS produces variety in the proteins translated from a limited number of genes through significant alterations in protein conformations that modulate cell functions (Yura *et al.* 2006). Our findings (Fig. 3; Table S3) indicate that alternative splicing may be an important mechanism by which the function of ASIP, *TYRP1*, *DCT*, and *RAB27A* are regulated in melanocytes. Therefore, it is necessary to identify the isoforms of these genes and their roles in pigmentation in the future. Interestingly, in the present study, we did not observe alternative splicing of the *MITF* gene in the skin, whereas multiple splice variants of *MITF* were previously found in other species such as sheep (Saravanaperumal *et al.* 2014), mice (Bismuth *et al.* 2005), and humans (Kuiper *et al.* 2004). There might be differences in the regulatory mechanisms of skin pigmentation between goats and the species mentioned above. Another valuable finding of the present study is the DE genes identified in the fetal skin between the two goat groups. In particular, members of the keratin family were highly differentially expressed in the fetal skin between the two goat groups, including *KRT1*, *KRT81*, *KRT83*, *KRTAP13-1*, *KRTAP11-1*, and *KRTAP7-1* (Table S5). Keratins make up the largest subgroup of intermediate filament proteins and represent the most abundant proteins in epithelial cells. These proteins are essential to sustain normal epidermal function and play a role in signaling (Porter and Lane 2003). Mutations of a member of the KRT family resulted in an unusual skin pigmentation in humans (Irvine *et al.* 1997; Horiguchi *et al.* 2005; Pascucci *et al.* 2006; Geller *et al.* 2013). Therefore, these DE keratins might participate in the signaling transduction involved in melanin synthesis in melanocytes or the transportation of melanin from melanocytes to keratinocytes through cellular interactions (Nakazawa *et al.* 1995; Seiberg 2001; Joshi *et al.* 2007).

Further functional clustering analysis of the DE genes showed that the terms involved in pigmentation and melanogenesis were highly overrepresented in our findings (Table S5 and S6), which is similar to the findings from studies on coat color in sheep (Fan *et al.* 2013) and skin color in the common carp (Jiang and Bikle 2014a). However, the specific DE genes of the skin differed in this study compared with the studies above. Specifically, the melanogenic *ASIP* (fold change >31) and *TYRP1* (fold change >87) were not only among the most significantly DE genes between the dark- and white-skinned goats, but were also associated with human pigmentation (Table 1). Previous studies have demonstrated that polymorphisms of *ASIP* are associated with darker skin color in African Americans (Bonilla *et al.* 2005) and fair skin color in Caucasians (Nan *et al.* 2009). *TYRP1*, a key regulator of melanin biosynthesis in melanocytes, is reported to be involved in the diversity of skin color in Europeans (Lao *et al.* 2007) and is a candidate gene for skin color (such as mouth, nose, and ear) in sheep (Raadsma *et al.* 2013). In addition, *DCT* is also associated with variations of skin pigmentation in Asians (Myles *et al.* 2007). However, the results of the present study suggest that *ASIP* is more likely the candidate gene for skin color in goats than *TYRP1* and *DCT*. We believe that *TYRP1* and *DCT* are two downstream genes whose expression levels are affected by *ASIP* in the KEGG pathway *melanogenesis* (Fig. 4B). However, efforts still be taken to ascertain the candidate genes underlying variations of skin color in goats using gene mapping techniques such as genome wide association studies (GWAS), next generation sequencing (NGS), or quantitative trait loci (QTL) analyses.

Our another valuable finding is the novel transcripts identified by RNA-seq (Table S1). Cabili, *et al* (2011) characterized a class of novel transcripts that were excluded by their coding potential criteria (a Pfam domain, a positive PhyloCSF score, or previously annotated as pseudogenes), and firstly termed them as TUCP (transcripts of uncertain coding potential) (Cabili *et al.* 2011). Since they merely focused the lincRNAs in human genome under a certain classification strategy, other subtypes of lncRNAs such as intronic lncRNAs and antisense lncRNAs were retained and thus grouped into the catalog of TUCP in their findings. However, in the present study, the intronic lncRNAs and antisense lncRNAs were absolutely excluded from the novel transcripts based on our recent study (Ren *et al.* 2016). Thus there are certain differences in classification of novel transcripts between the study from Cabili *et al* and ours. In view of subsequent analyses, we are certain that the identified transcripts with annotation and without annotation are two different groups of novel transcripts. Especially for those without annotation, the characteristics of a relatively low expression level and a highly enrichment of TE component (LINE/RTE-BovB) around TSS (transcription start site) (Fig 2) which are similar with that of the lincRNAs in recent studies (Cabili *et al.* 2011; Derrien *et al.* 2012; Kelley and Rinn 2012; Li *et al.* 2012; Nam and Bartel 2012; Billerey *et al.* 2014; Ren *et al.* 2016), strongly suggest this group of novel transcripts could be long nocoding RNAs. However, we are not certain that the novel transcripts with some annotation should be protein coding genes or long nocoding RNAs based on the current knowledge. It still need further efforts to identify their identity. Generally, this study provides a valuable resource for the genetic mechanisms involved in pigmentation diseases and contributes to the understanding of the biology of skin pigmentation and development in mammals.

## ACKNOWLEDGEMENTS

We thank Dr. Sun Wu, Sun Xiaowei, and their colleages from Southwest University for assistance in the sampling and experiments. We also thank Ms Yuan Jingxian and her colleagues for data processing in Novogene LTD. Co (Beijing). This work was supported by grants from the Chongqing Fund of application and development (cstc2013yykfC80003), the Chongqing Fundamental Research Funds (2013cstc-jbky-00106-zj; No.15435), and the Chongqing Fund of Agriculture Development (No.14412, No.15404).

## Additional files

Fig. S1. Classification of the raw reads by RNA-seq for each of the libraries (samples) in the hyperpigmented and normal fetal goats.

Fig. S2. Heatmap of the cluster analysis for the differentially expressed genes.

Table S1. Statistics of the mapping reads to the reference genome.

Table S2. Annotation of novel transcripts in Swiss-Prot database.

Table S3. Differential exon usage (DEU) analysis of the mRNAs in goat fetal skin.

Table S4. Differentially expressed genes between the wild-type and hyperpigmented goat skins.

Table S5. GO and KEGG analysis of the up-regulated DE genes using DAVID.

Table S6. Functional annotation clustering of the DE genes using DAVID.

